# Local Gradients of Functional Connectivity Enable Precise Fingerprinting of Infant Brains During Dynamic Development

**DOI:** 10.1101/2024.12.19.629222

**Authors:** Xinrui Yuan, Jiale Cheng, Dan Hu, Zhengwang Wu, Li Wang, Weili Lin, Gang Li

## Abstract

Brain functional connectivity patterns exhibit distinctive, individualized characteristics capable of distinguishing one individual from others, like fingerprint. Accurate and reliable depiction of individualized functional connectivity patterns during infancy is crucial for advancing our understanding of individual uniqueness and variability of the intrinsic functional architecture during dynamic early brain development, as well as its role in neurodevelopmental disorders. However, the highly dynamic and rapidly developing nature of the infant brain presents significant challenges in capturing robust and stable functional fingerprint, resulting in low accuracy in individual identification over ages during infancy using functional connectivity. Conventional methods rely on brain parcellations for computing inter-regional functional connections, which are sensitive to the chosen parcellation scheme and completely ignore important fine-grained, spatially detailed patterns in functional connectivity that encodes developmentally-invariant, subject-specific features critical for functional fingerprinting. To solve these issues, for the first time, we propose a novel method to leverage the high-resolution, vertex-level local gradient map of functional connectivity from resting-state functional MRI, which captures sharp changes and subject-specific rich information of functional connectivity patterns, to explore infant functional fingerprint. Leveraging a longitudinal dataset comprising 591 high-resolution resting-state functional MRI scans from 103 infants, our method demonstrates superior performance in infant individual identification across ages. Our method has unprecedentedly achieved 99% individual identification rates across three age-varied sub-datasets, with consistent and robust identification rates across different phase encoding directions, significantly outperforming atlas-based approaches with only around 70% accuracy. Further vertex-wise uniqueness and differential power analyses highlighted the discriminative identifiability of higher-order functional networks. Additionally, the local gradient-based functional fingerprints demonstrated reliable predictive capabilities for cognitive performance during infancy. These findings suggest the existence of unique individualized functional fingerprints during infancy and underscore the potential of local gradients of functional connectivity in capturing neurobiologically meaningful and fine-grained features of individualized characteristics for advancing normal and abnormal early brain development.

## Introduction

Resting-state functional magnetic resonance imaging (rs-fMRI) provides a promising and noninvasive imaging tool to measure the functional activities of the human brain in vivo and to map the individual functional connectivity (FC) between brain regions. Across various life stages, including infancy, adolescence, and adulthood, it has been observed that the human brain exhibits a unique and individualized FC pattern, akin to a “fingerprint”, which is remarkably robust to distinguish individuals from each other [1–4]. Since FC has been shown to remain highly stable over years after late adolescence [2, 3, 5, 6], it was utilized as the key approach to explore the existence of functional fingerprints, achieving high level of identifiability in most life stages. However, the identifiability of functional fingerprints in infants presents lower and less robust compared to relatively older age groups. This is likely due to the highly dynamic and complex maturation process occurring in the infant brain, which influences both its structure [7, 8] and function [9–12], These ongoing developmental changes result in evolving brain characteristics that are inherently more variable and difficult to interpret than those observed in more mature brains. Given that early developmental patterns during infancy play a critical role in shaping the subsequent cognitive and behavioral performances [5], accurately capturing these intricated and individualized brain functional characteristics is essential. Moreover, understanding the relationship between these early individualized functional patterns and behavioral phenotypes has significant potential for informing personalized diagnostic approaches and early intervention strategies aimed at optimizing developmental outcomes.

Traditionally, FC fingerprinting analyzes brain connectivity by mapping inter-region connections between a set of region-of-interests (ROIs) from predefined brain atlases, such as those containing 268 [1], 333 [13], 602 ROIs [14] or over 1000 ROIs [15, 16], which have been widely applied to reveal complex functional patterns in both mature and immature brains. In addition to atlas-based methods, numerous studies have employed principal gradient techniques, a vertex-wise approach, to capture global, brain-wide patterns and macroscale connectivity transitions along the cortical surface [17], as network topology has demonstrated significant behavioral implications [13, 18, 19]. By leveraging principal component analysis [20] and Laplacian eigenmaps [21], principal gradient-based approaches have successfully mapped functional organization and connectivity patterns across primary sensory and motor cortex as well as transmodal regions [22–24]. However, while atlas-based FC and vertex-wise principal-based approaches have the potential to identify individualized characteristics, they share similar limitation on merely capturing global-scale information. During infancy, the neurodevelopment is governed by dynamic processes that the functional brain networks were gradually subdivided with intra- and inter-modular connectivity, indicating the enhancement of functional segregation and integration [25, 26]. Under these rapid developmental conditions, it is challenging for global-scale FC to capture the subtle and stable “fingerprint”-like patterns that define early functional fingerprint. Moreover, reliance on predefined brain atlases imposes inherent constraints on connectivity analysis, particularly in developmental studies, as it not only limits the spatial resolution of analysis but also introduces difficulties related to cross-atlas comparability of statistical findings [27]. These limitations underscore the necessity for more sophisticated approaches capable of capturing intricate local spatial patterns that characterize the brain function development throughout infancy.

To address this issue, we adopt the local gradient of functional connectivity to investigate the individualized characteristics during infancy. Unlike the previous approaches, the local gradient methodology, widely applied in brain parcellation studies [13, 18, 19], is highly sensitive to detecting localized abrupt transitions in resting-state functional connectivity patterns across the cortical surface. This method captures different scales of architecture boundaries and is more attuned to certain biological boundaries than global-scale approaches. This capacity to detect spatially discrete abrupt changes across the cortical surface suggests the enhanced intra-individual stability of the local gradient approach during rapid developmental stages of infancy, making it particularly suitable for individualized parcellation and the follow-up phenotypes and behavior analysis for developing human brains [15]. As such, it offers distinct advantages for capturing nuanced individualized functional characteristics in developmental neuroscience research. To the best our knowledge, the application of local gradient approaches for brain functional fingerprinting remains unexplored. To mitigate this gap, in this work, we explore the potential to directly apply the local gradient method to characterize individualized characteristics of brain function, thereby facilitating the exploration of depicting complex and early individualized intervention strategies while eliminating the prerequisite step of individualized parcellation.

In general, the objective of this study is to delineate a vertex-wise infant functional connectivity fingerprint utilizing local gradient maps (LGMs) without reliance on any atlas. Specifically, our goals were to: (i) determine whether local gradient maps are capable of distinguishing individuals in capturing individualized functional characteristics during infancy; (ii) identify which functional networks manifest the most individualized uniqueness during infancy using local gradients of functional connectivity; (iii) examine the association between LGM-based functional fingerprint and early cognitive performance. Addressing these questions holds novel significance in neuroscience, yielding new perspectives on individualized intrinsic patterns of functional organization without reliance on atlas-based frameworks. Furthermore, it establishes a local-scale methodology that holds potential for characterizing individual-specific functional patterns, with implications for both phenotypic assessment and longitudinal monitoring of neurodevelopmental disorders during early brain development.

## Results

Fingerprinting analyses were performed on 103 individuals of the Baby Connectome Project (BCP) dataset [28] aged from 16 to 874 days. Study inclusion criteria required a minimum of two resting-state fMRI (rs-fMRI) scans meeting quality control standards. The cohort, comprising 103 subjects with 591 longitudinal rs-fMRI scans, was stratified into three distinct datasets (I, II, and III), characterized by varying age distributions and session gaps (the time interval between two fMRI sessions). Within each dataset, subjects contributed two longitudinal scans at different ages with session 1 (S1) earlier than session 2 (S2). Specifically, Datasets I, II, III are comprised of the first two sessions, the last two sessions, and the first and last sessions of each individual, with the mean session gaps were 147, 191, and 298 days, respectively. Detailed demographic information of these datasets is presented in **Figure 1**.

**Figure 1.**
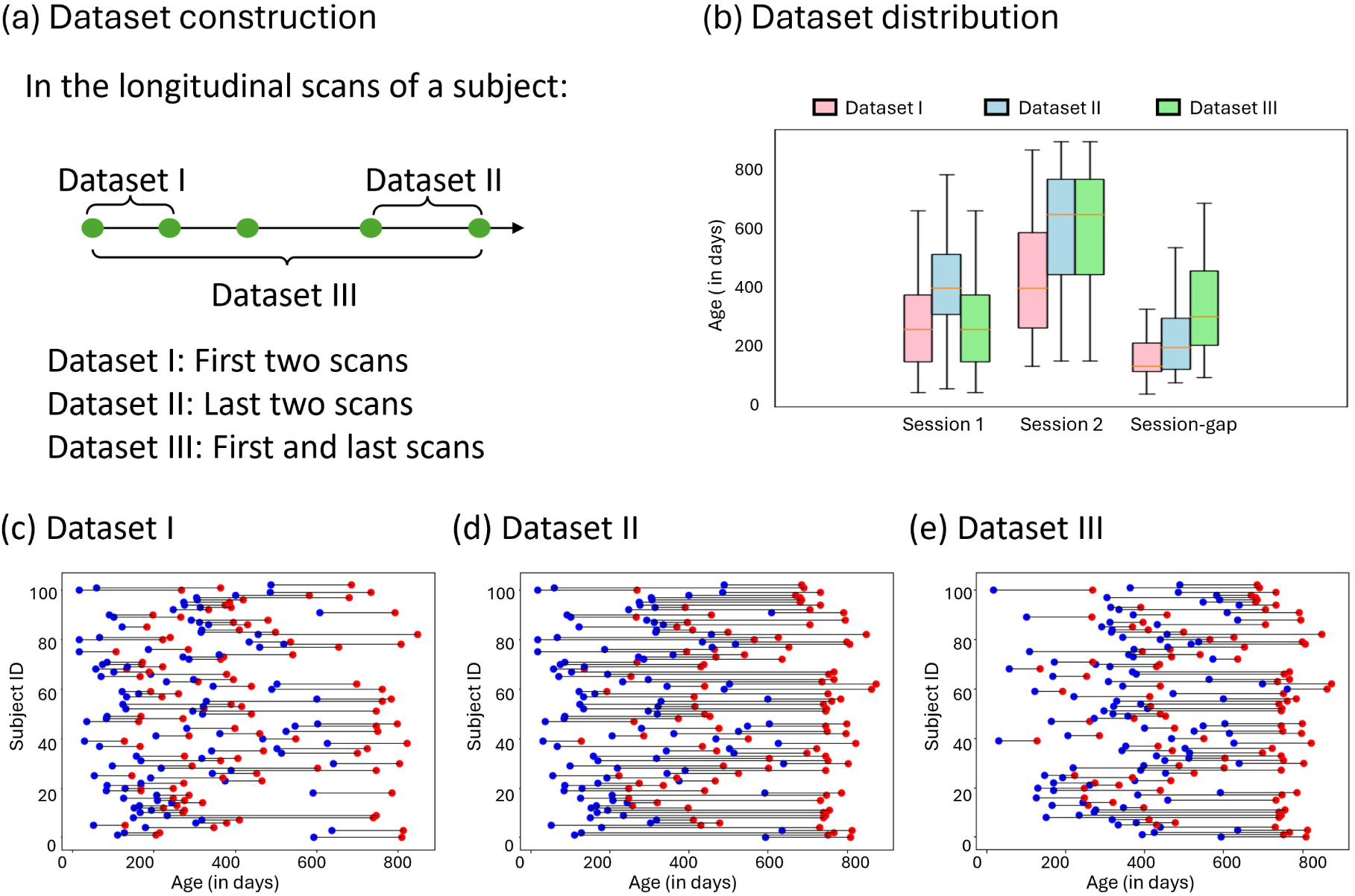
Dataset description. (a) Dataset construction. To study the LGM-based fingerprinting using different distributions of age and session gaps, Datasets I, II, and III consist of the first two scans, the last two scans, and the first and last scans for each subject, respectively. (b) The detailed age distribution information of the three datasets, the box plots of the scan age of session 1, session 2, and the session gap between session 1 and session 2. (c)-(e) The scan distribution of Datasets I–III, where the blue and red points represent the scans acquired from session 1 and session 2, respectively.

Local gradient maps (LGMs) of each rs-fMRI with 40,962 vertices were computed [15] after data processing (see Methods). Then the local correlation map (LCM) was constructed by computing Pearson’s correlation coefficients between each possible pair of patches centered at the same vertex on pairs of LGMs, while each vertex of LCM represents the similarity between two corresponding patches of two LGMs.

Identification was performed across pairs of sessions, each of which consists of one ‘target” and one ‘base’ session. If S1 is selected as the target session, then S2 will be the based session, and vice versa. In the following results, we selected S1 as the target session, and S2 as the base session unless we noticed specifically. For each individual’s LGM from the target set, it would be paired with each LGM in the base set to construct the LCM and build a set of LCMs. Notably, each vertex of the LCM represents the local similarity between pairs of LGMs. Then, with the set of LCMs, we compared the LCMs in a vertex-wise way, i.e., for each vertex, the LCM with the highest similarity value was regarded as the predicted identity at this vertex. Finally, through a majority voting across all vertices, the identity of the LCM which contributed the most the-highest-similarity-value vertices was considered as the final identified individual. The identification rate was measured as the percentage of subjects whose identity was correctly recognized out of the total number of subjects (see **Methods**).

For the Early Learning Composite (ELC) score prediction analysis, an additional 39 subjects with single scans were included with the previous 103 subjects to further enlarge the dataset.

### Local gradient map-based functional fingerprinting

#### Whole-brain identification

First, identification was performed using all vertices of LCMs (40,962 vertices). The identification rates were 103/103 (100%), 102/103 (99%), and 101/103 (98.1%) with S1 as target and S2 as base sessions in Dataset I, II, and III, respectively. The identification rates were 103/103 (100%), 102/103 (99%), and 101/103 (98.1%) with S2 as target and S1 as base sessions in Dataset I, II, and III, respectively. As the identical results showed in S1-S2 and reverse S1-S2 sessions, we only report the results in S1-S2 sessions unless we noticed specifically.

#### Network-based identification

We next tested identification accuracy on the basis of each of the ten specific functional networks to test whether certain brain networks contribute more to individual subject discriminability than others using LGM. Specifically, we adopted the networks definition based on a fine-grained functional parcellation maps of infants [10]. In this comparison, we predict the identity by calculating the-highest-similarity-value vertices in a certain network and use the identification rate to reflect the discriminability of this network for characterizing infants. We reported the average identification rates across three datasets for each functional network. Three networks emerged as the most successful in the individual identification: default mode network (network 4), posterior frontoparietal network (network 6), dorsal attention network (network 7), and anterior frontoparietal (network 9) with average identification rates across three datasets of 92.2% (93.2%, 93.2%, 90.3%), 90.3% (91.3%, 92.2, 87.4%), 90.3% (91.3%, 94.2%, 85.4%), and 90.3% (93.2%, 90.3%, 87.4%), respectively (**Figure 2b**).

**Figure 2.**
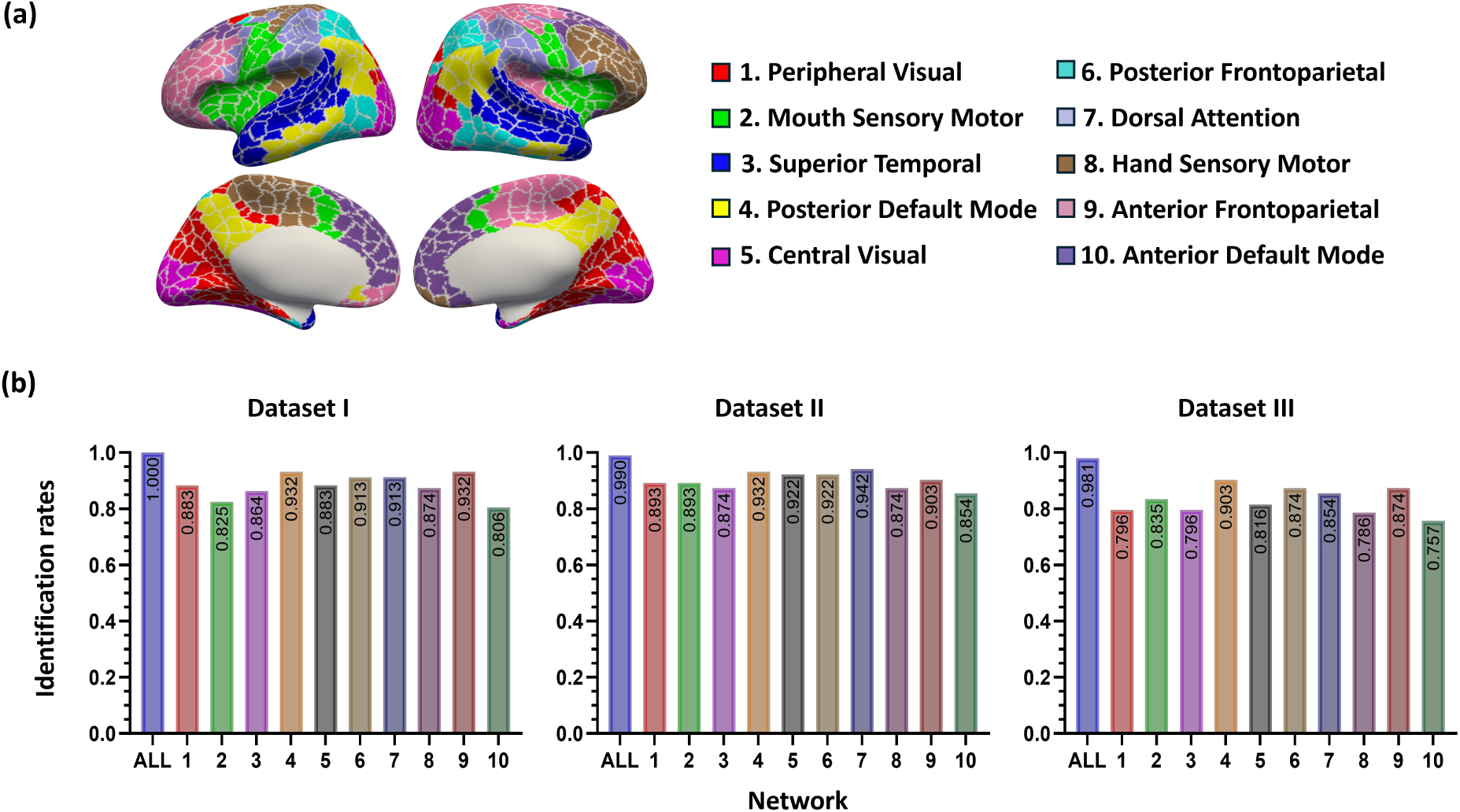
The network definition and identification rates across datasets and networks. (a) Network definition. We leveraged an infant-specific fine-grained brain functional parcellation map with 864 cortical region-of-interests (ROIs) grouped into ten networks [15]. (b) Identification rates based on whole-brain local gradient maps (ALL) and ten single networks.

### Factors affecting identification accuracy

#### Identification using different patch sizes and patch numbers

We conduct this experiment to investigate the impact of the resolution of local gradient maps on their ability to characterize the individualized infant functional characteristics. The computation of LCM is parameterized by two variables: patch number (the number of sampled vertices) and the patch size (defined by *r*-ring neighborhood) on the LGM. We firstly uniformly sampled vertices across the LGMs, followed by the selection of varying patch size centered on each sampled vertex. The patch size determines the spatial extent of local information incorporated into the similarity computation between different LGMs, with larger patch sizes incorporating more extensive local topological information.

We repeated the identification using different combinations of patch number and patch size, where patch number of LGM within {40,962, 10,242, 2,562} and patch size was based on the *r*-ring neighborhood (*r*∈{1, 2, 3, 4, 5, 6, 7}). The results (**Figure 3a**) demonstrated that the highest identification rate was achieved with patch number of 40,962 and patch size of 1-ring neighborhood.

**Figure 3.**
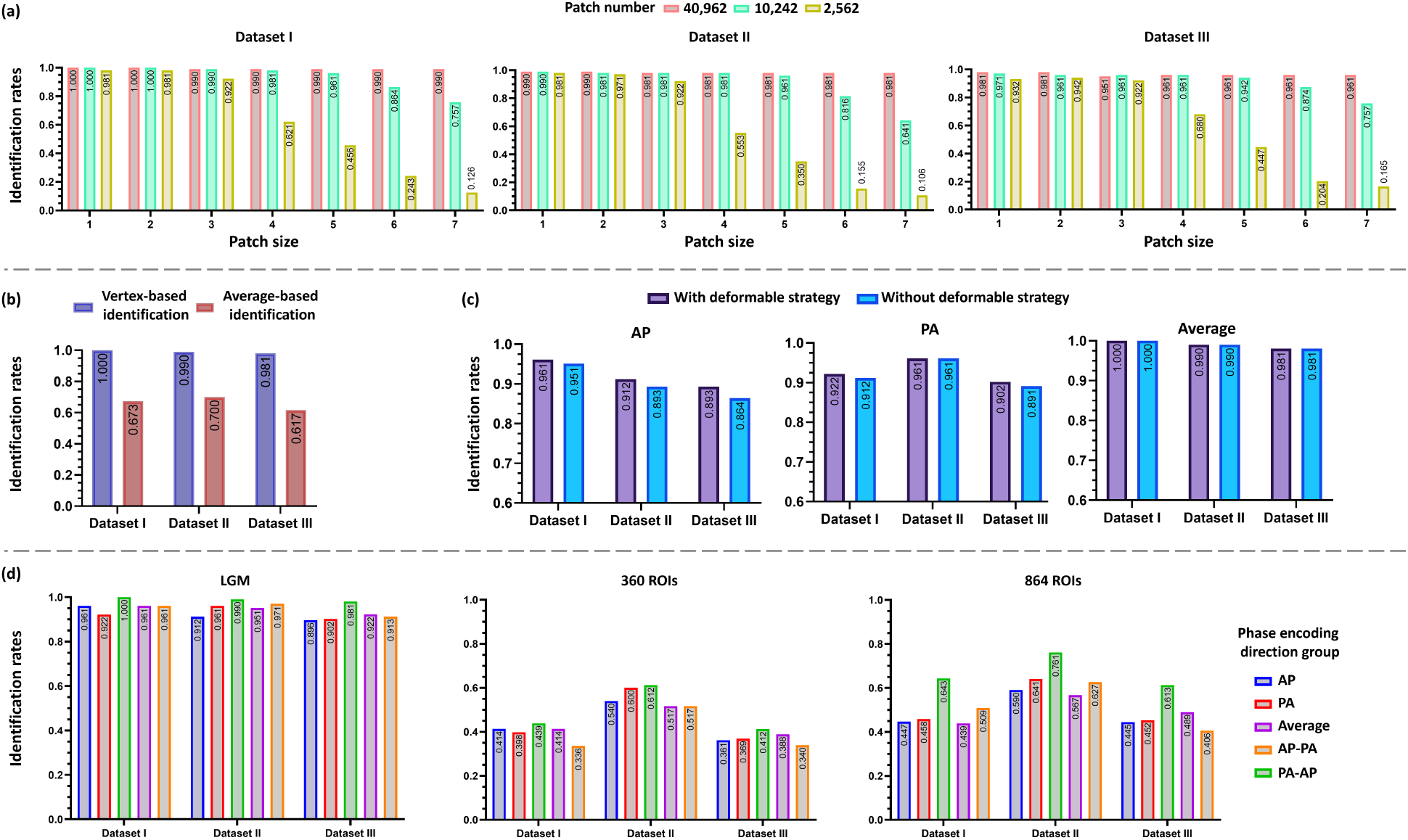
Factors affecting identification accuracy. (a) Identification using different patch sizes and patch numbers. (b) Identification based on different identification procedures. (c) Identification using deformable strategy. (d) Identification using different phase encoding directions via LGM and atlas-based functional connectome approaches (360 ROIs [29] and 864 ROIs [15]).

Specifically, at the highest resolution of 40,962 patches, Dataset I maintained consistent identification rates of 100% across all patch sizes. Dataset II demonstrated identification rates of 99% for the two smallest patch sizes, followed by consistent rates of 98.1% for larger sizes. Dataset III exhibited slightly lower but stable rates, ranging from 98.1% to 96.1% as patch size increased.

At an intermediate resolution of 10,242 patches, Dataset I showed a gradual decline in identification rates from 100% to 75.7% with increasing patch size. Similarly, Dataset II demonstrated a progressive decrease from 99% to 64.1%, while Dataset III exhibited rates declining from 97.1% to 75.7%.

At the lowest resolution of 2,562 patches, the impact of increasing patch size was most pronounced. Dataset I showed a marked decline in identification rates from 98.1% to 12.6%. Dataset II followed a similar pattern, decreasing from 98% to 10.7%, while Dataset III demonstrated a reduction from 93.2% to 16.5%. These results systematically demonstrate the inverse relationship between patch size and identification accuracy, particularly at lower resolutions.

Stating the effect of patch number, LCM computation based on the finer resolution (more patches) of the LGMs, the identification rates reach better performance in the three datasets; considering the effect the patch size, the increasing of patch size results in worse performance using finer resolution while results in better performance using coarser resolution (less patches), which indicating that the local changes and fine-grained information are more important and should be well captured in identification based on LGMs. Since the highest identification rate was achieved with patch number of 40,962 and patch size of 1-ring neighborhood, all the validations were based on this combination.

#### Identification based on identification procedures

The identification procedure employs an iterative methodology wherein each individual’s LGM from the target set is systematically compared against all LGMs in the base set to generate LCMs. To verify the significance of local subtle patterns, we explored two distinct prediction procedures for LCM comparison: vertex-wise identification and average-based identification. In the vertex-wise identification approach, which served as our primary identification method, all LCMs were initially combined to build an LCM set. Subsequently, vertex-wise comparison between the LCMs provides one recognized identify for each vertex, while the final predicted identity was determined through majority voting across all vertices. Conversely, the average-based identification method first computed the mean value of each LCM to obtain a single similarity metric, and then compared the single similarity metric among the set of LCMs to determine the predicted identity.

The vertex-based identification was outperformed than average-based identification, which neglects abundant fine-grained correlations resulting in inferior identification (**Figure 3b**). Specifically, identification rates of 103 infants using vertex-based identification were 100% (Dataset I), 99% (Dataset II), 98.1% (Dataset III), while 67.3% (Dataset I), 70% (Dataset II), 61.7% (Dataset III) using average-based identification. These results demonstrated that vertex-wise identification using LGMs has superior performance for capturing local and subtle information on the cerebral cortex.

#### Identification using different phase encoding directions

The majority of functional connectivity studies in the literature based on rs-fMRI data use either an anterior-to-posterior (AP) or a posterior-to-anterior (PA) phase encoding direction (PED). However, due to quality control requirements or participant availability constraints, paired AP and PA scans may not be consistently available for all sessions. To investigate the potential influence of PED on individualized functional characteristics derived from LGMs, we systematically evaluated functional fingerprinting across five experimental configurations: (1) both target and base sets comprising AP scans exclusively (AP group); (2) both target and base sets comprising PA scans exclusively (PA group); (3) target set comprising AP scans with base set comprising PA scans (AP-PA group); (4) target set comprising PA scans with base set comprising AP scans (PA-AP group); and (5) both target and base sets comprising averaged AP and PA scans (Average group). The LGM-based functional fingerprint demonstrated robust and consistent performance when target and base sets shared the same PED (**Figure 3d**). For the AP group, identification rates across Datasets I, II, and III were 96.1%, 91.2%, and 89.3%, respectively. Comparable performance was observed in the PA group, with rates of 92.2%, 96.1%, and 90.2%. Notably, the average group exhibited superior performance, achieving identification rates of 100%, 99%, and 98.1% across the three datasets.

When analyzing cross-PED configurations (AP-PA and PA-AP groups), though slightly lower, the identification rates maintained relative stability. The AP-PA group achieved rates of 96.1%, 95.1%, and 92.2% across Datasets I, II, and III, respectively, while the PA-AP group demonstrated rates of 96.1%, 97.1%, and 91.3%. These results suggest that the LGM-based approach maintains robust performance even across different phase encoding directions.

#### Identification using deformable strategy

To account for potential registration misalignment during data processing and enhance robustness against registration errors, we implemented a deformable strategy in the LCM computation (see **Methods**). This approach incorporated both patch rotation and patch translation, specifically allowing for patch rotations within a ±60° range and translations along six directional axes. We repeated the vertex-wise identification with deformable strategy (**Figure 3c**).

Implementation of the deformable strategy yielded notably high identification rates across datasets I-III: 96.1%, 91.2%, and 89.3% in the AP group; 92.2%, 96.1%, and 90.2% in the PA group; and 100%, 99%, and 98.1% in the averaged group, respectively. These results demonstrated improvement over the non-deformable LGM approach, which achieved rates of 95.1%, 89.3%, and 86.4% in Dataset I, II, and III for the AP group, and 91.2%, 96.1%, and 89.1% for the PA group. This enhancement suggests that the deformable strategy effectively mitigates local patch misalignment and reduces interference from signal noise and preprocessing artifacts, leading to more robust performance. Notably, the average group showed no additional improvement with the deformable strategy, suggesting that the averaging of AP and PA scans alone provides sufficient noise reduction, thereby diminishing the impact of the deformable approach in this context.

#### Comparison with connectome-based fingerprinting

To evaluate the relative robustness of individualized characteristics, we conducted a comparative analysis between LGM-based functional fingerprints and traditional connectome-based functional connectivity (FC) approaches across the previously described phase encoding directions. The connectomes were derived by utilizing two established whole-brain functional atlases [15, 29], incorporating 360 and 864 regions of interest (ROIs), respectively.

Connectome-based fingerprinting analysis was implemented across five PED configurations as mentioned above (**Figure 3d**). In the AP group, utilizing 360 ROIs yielded identification rates of 41.4%, 54%, and 36.1% for Datasets I, II, and III, respectively. Implementation with 864 ROIs demonstrated enhanced identification rates of 44.7%, 59 %, and 44.5% across the same datasets.

For the PA group, identification rates using the 360-ROI atlas were 39.8%, 60%, and 36.9% across Datasets I, II, and III, while the 864-ROI parcellation yielded improved rates of 45.8%, 64.1%, and 45.2% across the same datasets.

The average group exhibited identification rates of 43.9%, 61.2%, and 41.2% with 360-ROI atlas, and notably higher rates of 64.3%, 76.1%, and 61.3% with 864 ROIs across the three datasets.

In the AP-PA group, identification rates were 41.4%, 51.7%, and 39.8% using 360-ROI atlas, while the 864-ROI atlas achieved rates of 43.9%, 55.7%, and 48.9% across Datasets I-III.

Similarly, the PA-AP group demonstrated identification rates of 33.6%, 51.7%, and 34% with 360 ROIs, and 50.9%, 62.7%, and 40.6% with 864 ROIs across the three datasets.

These findings indicate that connectome-based fingerprinting exhibits sensitivity to PED conditions, necessitating consistent data acquisition conditions to achieve satisfactory identification rates. The performance of connectome-based methods remains inferior to that achieved through LGM-based functional fingerprinting.

#### Quantifying vertex-wise contributions to identification

To quantify the extent to which different vertices contribute to identity identification and capture the most individualized functional characteristics, we derived two measures: uniqueness (𝕌) and differential power (DP) [1, 30–32]. Uniqueness was highest for vertices that exhibited substantial variability among individuals while maintaining consistency across MRI sessions within the same individual. DP, on the other hand, reflects each vertex’s ability to distinguish an individual by quantifying how “characteristic” that vertex tends to be. Vertices with high DP values demonstrate consistent measurements within an individual across MRI sessions but show significant differences across individuals, regardless of condition. In other words, vertices with high DP values are thought to make a substantial contribution to individual identification (see **Methods**).

We calculated the average vertex-wise DP and ***U*** for all vertices across the three datasets and computed the network-wise DP and ***U*** to determine which network has significant contribution to identification (**Figure 4**). First, our analysis revealed that vertices with high uniqueness values were primarily located in the occipital lobe, superior frontal gyrus, and were associated with the central visual, posterior frontoparietal, and posterior default mode networks. These vertices demonstrated high consistency within the same individuals while varying across other individuals. Similarly, vertices with high DP values were predominantly found in the superior frontal gyrus, indicating that these vertices are highly discriminative and play a critical role in individual identification.

**Figure 4.**
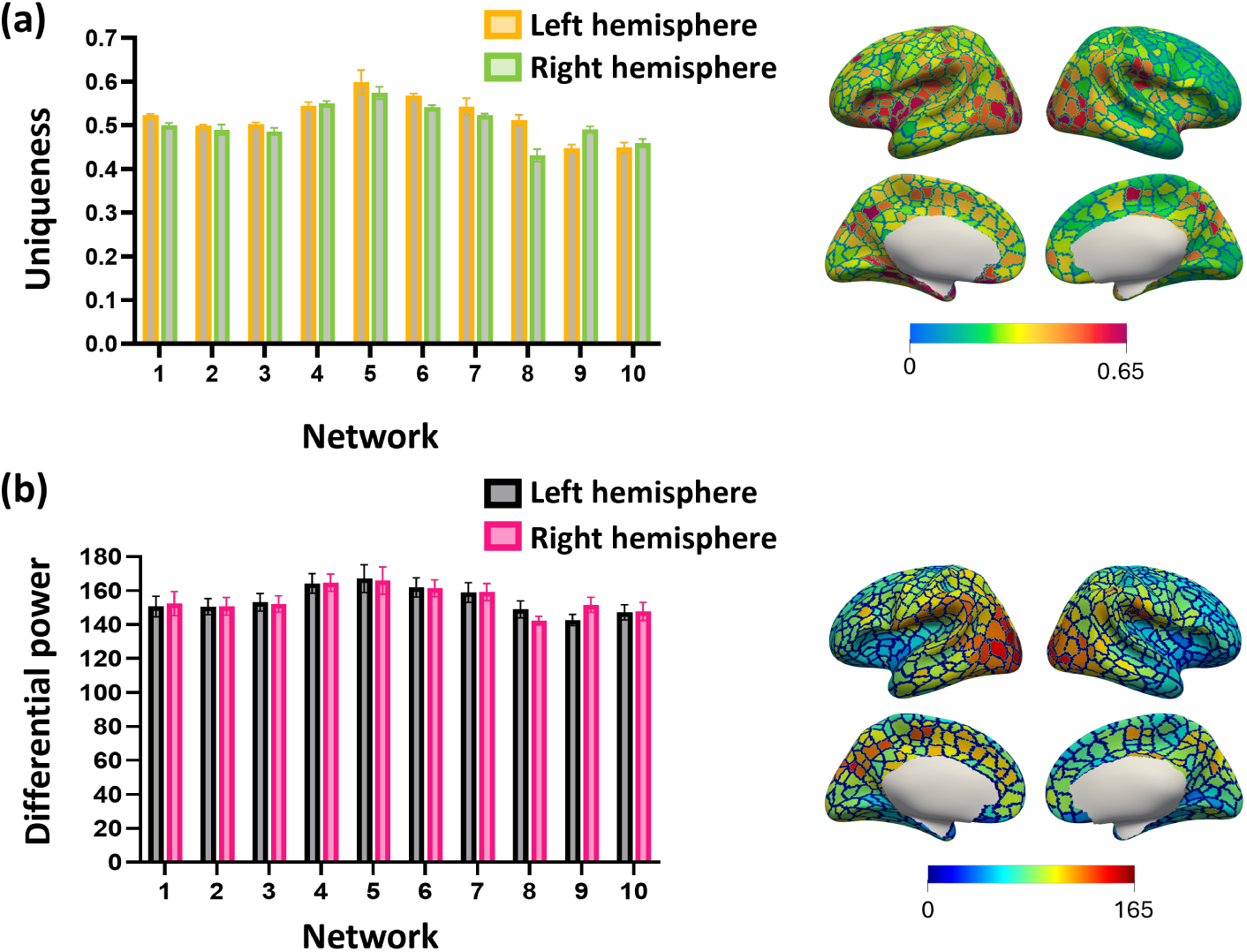
Vertex-wise contributions to identification. (a) Network-wise average uniqueness map. The color of the bars, yellow/green in the top row and black/magenta in the bottom row, indicates left and right hemispheres, respectively. (b) Network-wise average differential power map. The bars represent the mean and std. of the corresponding values across the dataset I, II, and III.

### Cognitive Score Prediction

Relatively few studies have explored LGM-based features in infant healthy participants or investigated its association with cognitive performance. In this study, we replicated the cognitive prediction following [1, 14] to validate whether such an association exists in LGM-based functional features and cognition. We found LGM produced significant predictions, with an *r*-value of 0.244 and a *p*-value smaller than 0.05.

To evaluate the differential contributions of functional networks to cognitive prediction, we implemented a network-wise analysis. Within the framework of 10 iterations 10cross validation, the selected vertices for prediction varied slightly across each fold. We computed the mean number of selected vertices per functional network, normalized by the total vertex count within each network to account for network size heterogeneity (**Figure 5b**). Our analysis revealed that the visual system, dorsal attention, and anterior frontoparietal networks made predominant contributions, particularly in terms of positively correlated vertices. These findings suggest a pivotal role in visual functional areas in local gradients of functional connectivity analyses.

**Figure 5.**
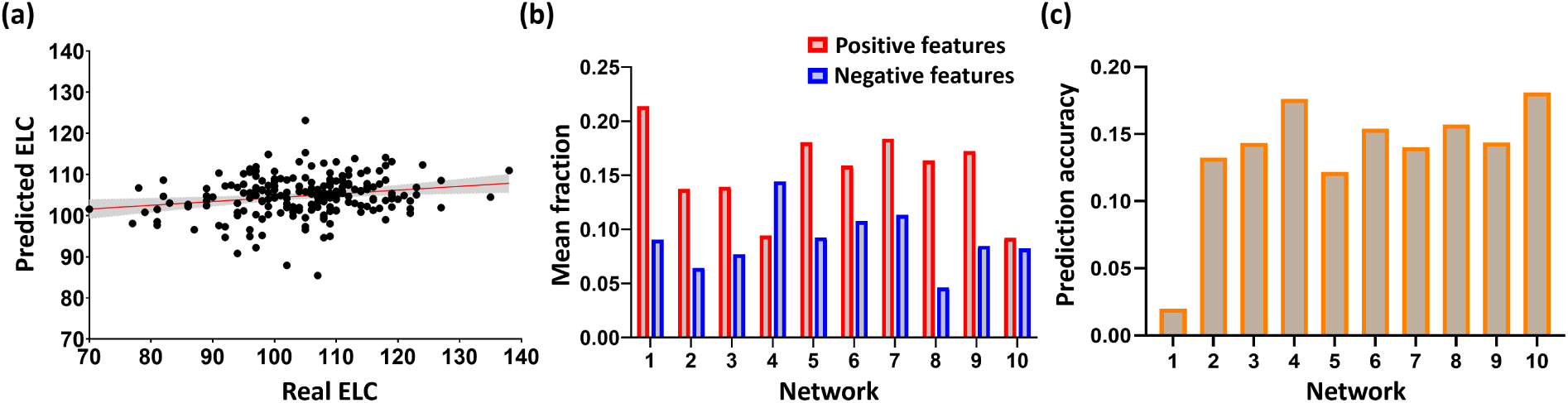
Cognitive score prediction based on local gradient maps (LGMs). (a) ELC score prediction based on LGM of whole brain. The scatter plot shows the predicted ELC from one time 10-fold cross-validation and the real observed ELC. Each dot represents the predicted and real ELC pair from one subject; the gray area indicates 95% confidence interval for the fit line. (b) Mean fraction of within-network vertices selected in cognitive prediction, positive- and negative-feature represented by red and blue, respectively. Y axis indicates the mean fraction of vertices selected across all repeated iterations; X axis indicates network labels as illustrated in Figure 2a. (c) The prediction accuracy (*r*-value, Pearson’s correlation between predicted and real ELC scores) of leave-one-network-out strategy. Lower value indicates significant contribution of this network in cognitive prediction. Y axis indicates the *r*-value, while X axis denotes network labels as illustrated in Figure 2a.

To further validate the relative importance of single networks, we employed an iteratively leave-one-network-out approach, where cognitive scores were predicted with the remaining brain networks configuration, and the results were averaged across cross-validation results. Network importance was inversely proportional to the prediction accuracy (*r*-value, Pearson’s correlation between predicted and real ELC scores) of the remaining networks following its removal. Lower value indicates significant contribution of this network in cognitive prediction. This analysis corroborated our initial findings: the visual, dorsal attention, and anterior frontoparietal networks yielded the lowest prediction accuracy upon removal, thereby confirming their significant contributions of cognition prediction (**Figure 5c**).

## Discussion

In this study, we explored the capability of local gradient maps (LGM) of functional connectivity to describe individualized characteristics by applying them to fingerprinting and cognitive prediction across three datasets spanning different age ranges. Stable and accurate individual identification was achieved across varying LGM resolutions, identification procedures, and phase encoding directions (PED) in rs-fMRI acquisition. Notably, LGM consistently outperformed conventional atlas-based FC in all comparisons. Analysis of the most contributive vertices for identification, with uniqueness and DP maps, revealed that LGMs captures significant discriminative information primarily within posterior default mode network, posterior frontoparietal network, and dorsal attention network. Meanwhile, cognitive prediction analyses showed that regions associated with cognition, as identified through LGM, were predominantly located in visual network, dorsal attention network, and anterior frontoparietal network. Collectively, these findings highlight the presence of vertex-wise local individualized functional patterns embedded in LGMs, offering new insights into the characterization of individual traits and behavior phenotypes, particularly during infancy.

### Local gradient map-based functional fingerprint is of robust and accurate individualized functional characteristics

Our findings demonstrate that local gradients of functional connectivity exhibit distinct individual fingerprinting characteristics during early infancy, achieving significant greater accuracy and stability compared to traditional atlas-based original FC methods. Despite the rapid brain development during infancy, we achieved an average identification rate exceeding 98% with an average session gap approaching over 200 days. This marks a substantial improvement over previous studies, showing the existence of functional connectome fingerprints in developing brains, with reported identification rates of 11% in the perinatal period [33], 62% during infancy [14], and 42% in older children [34] using resting-state functional connectomes. A potential factor contributing to this significant improvement could be the metric used, i.e., LGM. Given the atlas-based FC yields to atlas selection and capture global information between regions, it may be less effective at capturing local individualized characteristics during the dynamic developmental stage [35].

While LGM demonstrates considerable reliability and robustness during early infancy, multiple factors likely contribute to the capability to capture the individualized characteristics. Our analysis revealed that vertex-based identification significantly outperformed the global average-based identification approach (**Figure 7a**), substantiating the crucial role of vertex-wise analysis in achieving high identification accuracy. This finding suggests that local similarity patterns between patches preserve richer information than global averaging approaches across the whole brain, which treat whole-brain connectivity as a single “big” patch, thereby fail to capture intra-regional relationships, same as the limitation inherent in connectome-based identification methods. Furthermore, we evaluated various combinations of patch number and patch size in LCM computation to assess how patch-wise similarity within LGMs affect the description of functional fingerprinting. At higher resolutions (e.g., 40,962 patches), patch size demonstrated minimal impact on identification performance. However, at lower resolutions (e.g., 2,562 patches), we observed an inverse relationship between patch size and performance, with larger patches leading to diminished performance. These results suggest that fine-grained patch sizes are essential for effectively capturing local similarity and dissimilarity patterns, emphasizing the importance of local spatial information in delineating intra-individual stability within functional connectivity gradients. Another additional factor is the influence of phase encoding direction (PED) in resting-state fMRI acquisitions, as varying PEDs can introduce distinct patterns of signal distortion and loss [38, 73]. Notably, our LGM-based functional fingerprint maintained significant identification rates with mixed PED scans compared to atlas-based approaches (**Figure 3d**). Atlas-based approach exhibited reduced accuracy with mixed PED scans compared to single PED acquisitions, suggesting its sensitivity to signal noise introducing by varied PEDs, while our local gradient map-based approach exhibited robust performance. Importantly, the identification rates improved when averaging local gradient maps from anterior-posterior and posterior-anterior acquisitions, consistent with findings observed in adult populations [36]. Additionally, LGM demonstrated the highest identification performances in Dataset I, followed by Dataset II and III, suggesting that LGMs are most effective during period of rapid brain development and when session intervals are short, as they capture more accurate and dynamic functional characteristics during this critical period.

Understanding how local gradients of functional connectivity capture unique and stable characteristics during brain development is a complex challenge. Prior research has shown that functional brain networks undergo progressive subdivision, characterized by evolving intra- and inter-modular connectivity patterns, indicating enhanced functional segregation and integration during development [25]. This developmental trajectory implies continuous modification of functional networks throughout infancy. The constraints imposed by predefined parcellations in connectome-based fingerprinting methodologies may inadequately account for these dynamical developmental changes, potentially explaining the reduced identifiability observed in longitudinal studies [14, 33]. Local gradients, on the other hand, have been demonstrated to reflect distinct aspects of architectonic boundaries and exhibit heightened sensitivity to specific biological demarcations, correlating meaningfully with behavioral outcomes [18, 19]. Notably, short-range connectivity develops until approximately the fourth month after birth [37, 38], preceding the maturation of long-range connections and reflecting individual characteristics in functional architecture [39, 40]. This highlights the utility of exploring individualized functional characteristics at a local level during this developmental period of infancy. These empirical observations offer a mechanistic explanation for the existence and stability of local gradients based functional fingerprint throughout infancy, attributable to their capacity to capture fine-grained spatial information that typically remains undetectable at global scales of analysis.

### Distinguishing cortical loci of local gradients of functional connectivity

Brain development follows a posterior-to-anterior maturational trajectory [41], wherein sensory systems maturing earlier than higher-order systems [42]. Notably, analyses of structural connectome data have revealed reduced identification rates when examining regions in isolation, indicating that whole-brain analyses provide more robust fingerprinting characteristics in structural connectivity [33]. This contrasts with functional connectivity analyses, where frontoparietal networks demonstrated superior identification rates compared to whole-brain measures [1, 14]. However, by utilizing LGM, while still origin from functional connectivity, the identification performance based on individual networks proved less effective compared to the whole-brain. This observation indicates that examining isolated network disruptions may be insufficient for capturing the full extent of local gradients of functional connectivity, emphasizing the importance of considering global context in LGM-based analyses.

Our analyses revealed peak identification rates in the posterior default mode network, posterior frontoparietal network, and dorsal attention network. These findings align with previous research demonstrating that higher-order systems contribute substantially to individual identifiability [1, 14]. LGM demonstrated robust identification performance in association networks, which exhibit distinct individual-specific characteristics. This finding suggests that LGM effectively captures subtle variations in functional higher-order patterns and detects localized changes that reflect individual-specific network organization. The method’s sensitivity to local topological differences, which are particularly pronounced in higher-order systems, underscores its utility in characterizing individual neural signatures. Notably, these results stand in contrast to observations in preterm neonates, where sensorimotor systems demonstrated superior identification rates [33]. This discrepancy may stem from differences in population characteristics, as functional cores within in high order networks appear largely incomplete and fragmented [43, 44].

### Association exists between local gradients of functional connectivity and cognitive behavior

We also demonstrated an association between LGM and cognitive behavior, consistent with exiting studies demonstrating the relationship between individual uniqueness and cognitive behavior [1, 2, 5, 14, 45]. Among the ten functional networks [15], our analyses identified the visual network, dorsal attention network, and anterior frontoparietal network as the principal predictors of cognitive function. The visual network, in particular, is relatively mature with adult-like topology after birth [26, 46, 47], with significant increases in functional connectivity strength and heterogeneity during early development [48]. These developmental patterns, which are closely linked to brain activity in cognitive behavior, can be easily captured by LGM at a local level.

Interestingly, previous research comparing LGM- and atlas-based approaches suggested that local gradient methodologies might lack behaviorally-relevant information [49]. This apparent discrepancy with our findings can be attributed to methodological distinctions in local gradient implementations. Our study employed an infant-specific local gradient methodology [15], which offers enhanced resolution of patterns in temporo-occipital, parietal, and lateral prefrontal regions compared to the conventional local gradient approach utilized in prior studies [13, 50]. These fundamental differences in gradient computation render direct comparisons of cognitive prediction results challenging. Additionally, the previous study’s use of kernel regression for behavioral prediction, without feature selection optimization, may have reduced performance by introducing irrelevant or noisy predictor variables.

### Additional Considerations

Our findings should be interpreted in light of several limitations. First, we focused on static functional connectivity to investigate vertex-wise individualized patterns. Exploring whether dynamic vertex-wise functional connectivity and dynamic measures [32] can characterize individual uniqueness would further improve our understanding of the inherent traits and ongoing changes. Second, all the experiments were conducted during infancy. Given the strong performance of local gradient-based functional fingerprints, it will be interesting to test their reproductivity in other age groups, e.g., prenatal, children, adolescence, and so on. Third, it has been shown that the principal gradients have extraordinary reproducibility across datasets and are relevant to behavior [49, 51, 52]. Future work should further explore these aspects to deepen our understanding of functional gradient-based fingerprints.

## Conclusion

Collectively, our findings reveal that LGMs capture abundant local information and subtle changes, enabling precise and stable functional fingerprinting during early infancy, despite the rapid early development occurring during this period. Additionally, we also found that fingerprints derived from LGMs outperformed the atlas-based counterparts. The stability and reliability of LGMs hold great promises for advancing local-level analyses in future neuroimaging studies.

## Methods

### Participants and imaging acquisition

Subjects in this study are from the UNC/UMN Baby Connectome Project (BCP) [28]. For the study of normal early brain development, all infants recruited in BCP were born at the gestational age of 37-42 weeks and were free of any major pregnancy and delivery complications. We used 103 subjects with 591 longitudinal scans acquired at different ages ranging from 16 to 874 days after term birth. All infant MRIs were acquired during infants’ natural sleep using a 3T Siemens Prisma MRI scanner with a Siemens 32-channel head coil. T1-weighted images (208 sagittal slices) were obtained by using the three-dimensional magnetization-prepared rapid gradient echo sequence: repetition time (TR), echo time (TE), and inversion time (TI) = 2400, 2.24, 1600 ms, respectively; flip angle = 8°, and resolution = 0.8 × 0.8 × 0.8 mm^3^. T2-weighted images (208 sagittal slices) with turbo spin-echo sequences (turbo factor = 314, echo train length = 1166 ms): TR, TE = 3200, 564 ms, respectively; and resolution = 0.8 × 0.8 × 0.8 mm^3^ using a variable flip angle. All structural MRI data were assessed visually for excessive motion, insufficient coverage, and/or ghosting to ensure sufficient image quality for processing. For the same cohort, rs-fMRI scans were also acquired using a blood oxygenation level-dependent (BOLD) contrast-sensitive gradient echo-planar sequence: TR = 800 ms, TE = 37 ms, flip angle = 80°, field of view = 208 × 208 mm, 72 axial slices per volume, resolution = 2 × 2 × 2mm^3^, total volumes = 420. fMRI scans include anterior to posterior (AP) scans and posterior to anterior (PA) scans, which are two opposite phase encoding directions (PEDs) for better correction of geometric distortions. Here, in our study, we tested our model in three groups: AP, PA, and average, where the average group combines the scans acquired from the same subject at the same age (averaging AP and PA scans). Besides, we used cognitive data that were collected within one month after each fMRI data collection. We used 5 cognitive scales consisting of expressive language, fine motor, receptive language, visual reception, and gross motor scores and ELC cognitive scores defined by [53, 54], which is a composite of the first four scores. We selected standard scores for our study. Additionally, a total of 39 scans from the BCP dataset were accompanied by concurrent behavior evaluation using the MSEL assessment, which included the early learning composite score and five sub-domains: gross motor, fine motor, expressive language, receptive language, and visual reception.

### Data processing

All structural and functional MRIs were processed following state-of-the-art infant-tailored pipelines [11, 55]. All reconstructed cortical surfaces were first registered onto the UNC 4D infant cortical surface atlas [56, 57] based on cortical geometric features. The functional time series at each vertex on the resampled cortical surface were then extracted. For each cortical surface, the local gradient map (LGM) of functional connectivity was then computed according to [15, 50] and further resampled with 40,962 vertices (**Figure 6**).

**Figure 6.**
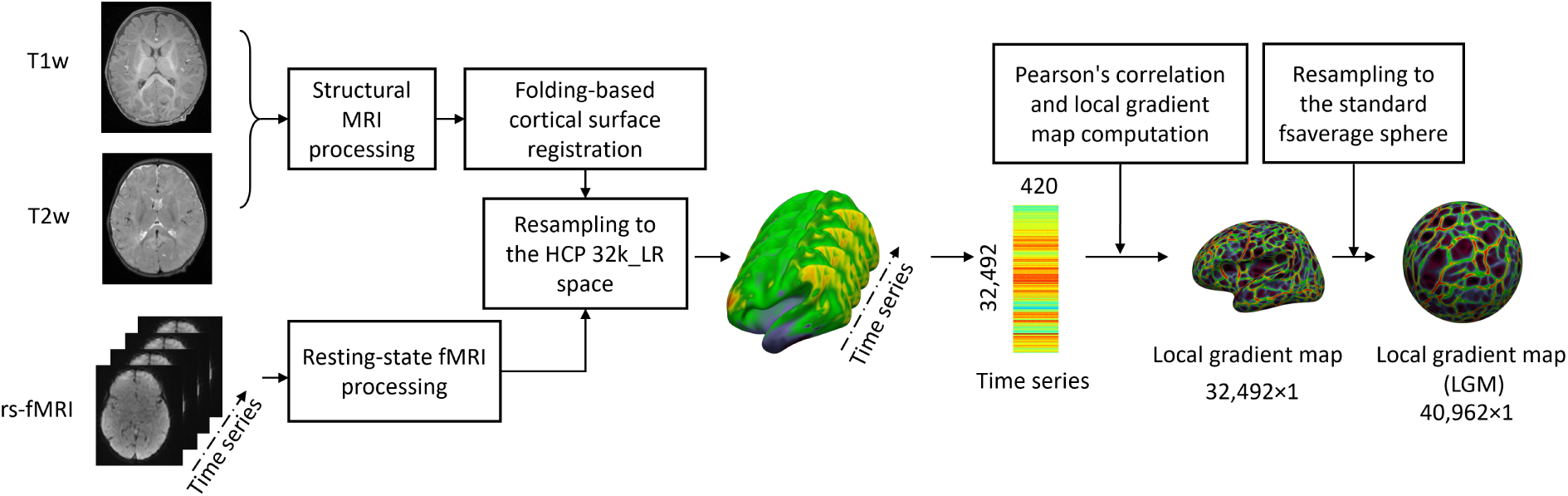
Data processing. All structural and functional MRIs were processed following state-of-the-art infant-tailored pipelines [11, 55]. All reconstructed cortical surfaces were first registered onto the UNC 4D infant cortical surface atlas [56, 57] based on cortical geometric features. The functional time series at each vertex on the resampled cortical surface were then extracted. The local gradient map (LGM) of functional connectivity was then computed according to [15, 50] and further resampled to 40,962 vertices.

#### Structural MRI processing

All T1- and T2-weighted magnetic resonance images underwent processing using an infant-specific pipeline [58] that has been extensively validated across numerous infant studies. The processing protocol encompasses multiple sequential steps: First, T2-weighted images were rigidly aligned to their corresponding T1-weighted images using FLIRT in FSL [59–61]. Subsequently, skull stripping was performed using a deep learning-based methodology [62], followed by manual refinement to ensure complete removal of skull and dura matter. The cerebellum and brain stem were then removed through atlas-based registration, followed by correction of intensity inhomogeneity using the N3 method [63]. Brain images underwent longitudinally consistent segmentation into white matter, gray matter, and cerebrospinal fluid utilizing an infant-dedicated deep learning-based approach [64]. Finally, each brain was separated into left and right hemispheres with subsequent filling of non-cortical structures.

#### Resting-state functional MRI processing

Resting-state fMRI data were preprocessed using an infant-specific functional pipeline in the following steps [11, 65]: correction of head motion and spatial distortion using FSL [59], registration of the rs-fMRI scans onto the T1w structural MRI using a boundary-based registration approach [66, 67], transformation and deformation fields were combined and used to resample the rs-fMRI data through a one-time resampling strategy, then the conservative high-pass filter with a sigma of 1000s was applied to remove the linear trends. After that, the individual independent component analysis was conducted to decompose each of the preprocessed rs-fMRI data into 150 components using MELODIC [68] in FSL and then an automatic deep learning-based noise-related component identification algorithm was used to identify and remove non-signal components to clean the rs-fMRI data [69].

#### Cortical surface reconstruction and mapping

Following tissue segmentation, inner, middle, and outer cortical surfaces were reconstructed using topology-preserving deformable surface methods [70, 71]. After the surface reconstruction, the inner cortical surface, which has vertex-to-vertex correspondences with the middle and outer cortical surfaces, was further smoothed, inflated, and mapped onto a standard sphere [72]. To ensure the accuracy of longitudinal analysis during infancy, it is necessary to perform longitudinally consistent cortical surface registration [57]. The process incorporates critical steps for longitudinal analysis: generation of intra-subject and population-mean surface maps, co-registration of individual surfaces using Spherical Demons [73], and mapping to the HCP 164k fs_LR space. All surfaces were ultimately warped and resampled to 32k fs_LR space [74] to establish vertex-to-vertex correspondence across subjects and ages. Finally, rs-fMRI time courses were resampled onto the middle cortical surface and spatially smoothed, creating a standardized framework for subsequent analysis.

#### Computation of individual local gradient map

We computed the LGM which identifies sharp transitions in resting-state functional connectivity (RSFC), inherently delineating boundaries between functional parcels. This approach has been extensively validated for generating meaningful fMRI-based cortical parcellations in adult populations [13, 18]. Our methodology processes anterior-posterior (AP) and posterior-anterior (PA) scans independently, integrating them only in the final step. For each scan, we construct a functional connectivity matrix by computing pair-wise correlations of BOLD signals between all cortical vertices in the CIFTI file, yielding a 32k × 64k RSFC matrix per hemisphere, with each row representing a vertex-specific RSFC map. The FC matrices then undergo Fisher’s r-to-z transformation, normalizing features across vertices to ensure comparative scaling. Subsequently, we generate a second-order correlation matrix (RSFC-2nd) sized 32k × 32k per hemisphere by correlating z-transformed RSFC maps across all cortical vertices within the same hemisphere. While primary RSFC maps exhibit gradual transitions, these second-order correlation maps demonstrate more pronounced transitions, thereby facilitating the identification of functional boundaries between adjacent areas [19]. Following the methodology of [29], we calculate the functional gradient on the RSFC-2nd, producing a 32k × 32k gradient matrix per hemisphere. Utilizing watershed-based boundary detection protocols [13], we generate 32k binary boundary maps per hemisphere from the gradient matrix. The local gradient map is defined as the mean of the 32k binary boundary maps. For scan sessions containing both AP and PA acquisitions, the local gradient maps from AP and PA scans are averaged for each session. Finally, the local gradient maps were further warped to the fsaverage space for further computation.

### Local gradient map-based identification test

The vertex-wise LGM-based identification test consists of one target set, 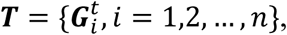 and one base set, 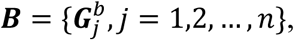 where *n* denotes the number of subjects, 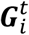 and 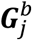 are LGMs at session *t* and session *b*, respectively. In the process of identification, one target 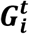 was selected from target set ***T***, and Pearson’s correlation coefficient was used to measure the vertex-wise similarity map between 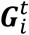 and all LGMs in base set ***B***, where the similarity map was named as local correlation maps (LCM).

#### Computation of individual local correlation maps

For each sampled vertex *v* in local gradient map (LGM), we first uniformly sampled its local surface patches, 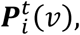 where *v* = 1,2, …, *V*, of the LGM with *r*-ring neighborhood for representing their spatially detailed functional patterns, and the local correlation map (LCM) is computed between corresponding patches of two LGMs, and this procedure is defined as: 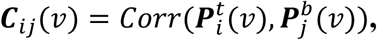 where *Corr* is Pearson’s correlation coefficient (**Figure 7a**).

**Figure 7.**
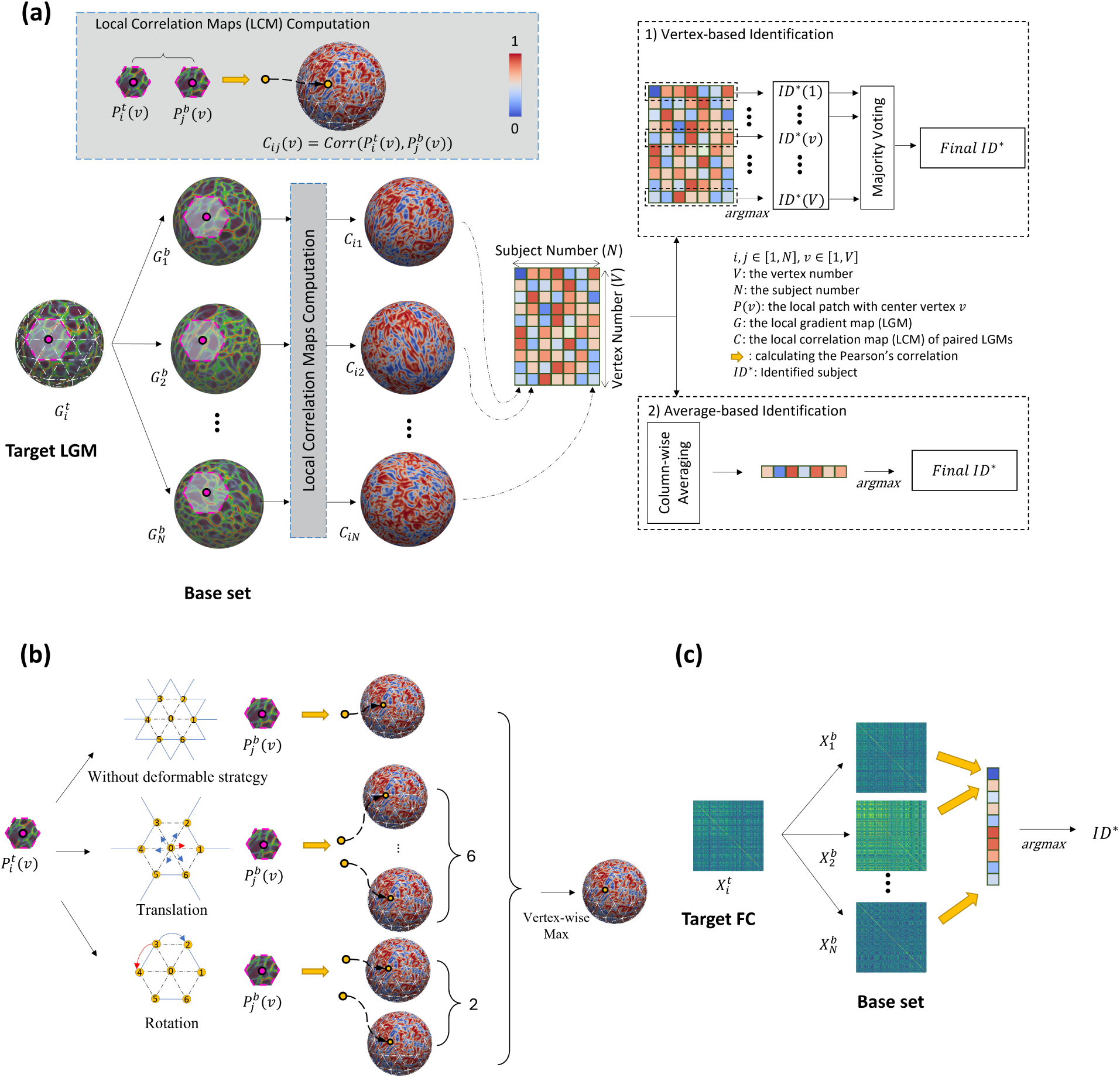
The process of identification using local gradient maps (LGMs) and connectome-based approaches. (a) The identification process of local gradient maps. The vertex-wise LGM-based identification test consists of one target set, 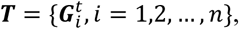 and one base set, 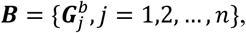 where *n* denotes the number of subjects, 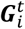 and 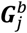 are LGMs at session *t* and session *b*, respectively. For each target 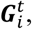 Pearson’s correlation coefficient was used to measure the vertex-wise similarity map between 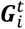 and all LGMs in base set ***B***, where the similarity map was named as local correlation maps (LCM). After computing all LCMs iteratively for all subjects in ***T***, we concatenated the *n* LCMs and formed a local correlation matrix, where each column represents an LCM of a pair of LGMs. For vertex-based identification solution, the identified subject was determined by the highest similarity at each vertex, then by the majority voting across vertices, we can determine the final identity. Conversely, the average-based identification method first computed the mean value of each LCM to obtain a single similarity metric, and then compared the single similarity metric among the set of LCMs to determine the predicted identity. (b) The details of computation of local correlation maps (LCMs) using deformable patch strategy. This approach incorporated both patch rotation and patch translation, specifically allowing for patch rotations within a ±60° range and translations along six directional axes of LGMs, which generates additional 6 and 2 LCMs, respectively. By combining the original LCM, two LCMs with rotation, and six LCMs with translation, the maximum value at each vertex was determined the final LCM. (c) The identification process of using atlas-based functional connectome. The atlas-based individual infant identification test was performed across paired scans consisting of one from target set, 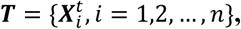 and one from base set, 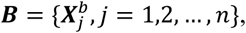 where 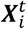 and 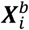 are all flattened vectors of the upper triangle of the whole-brain connectivity matrix and the subscripts ***i*** and ***j*** denote the subject index. In the process of identification, one target matrix, 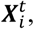 was selected from the target set ***T***, and the Pearson’s correlation coefficient was used to measure the similarity between 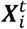 and all matrices in base set ***B***. The identity in ***B*** having the largest similarity with 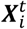 was regarded as the identified subject.

There are different alternatives for resolution of LCMs. Specifically, we computed LCMs with sampled vertex number of LGM within {40,962, 10,242, 2,562} and patch size was based on the *r*-ring neighborhood (*r*∈{1,2,3,4,5,6,7}). Of note, the sampled vertices were uniformly distributed on the standard fsaverage sphere.

Additionally, to get rid of the potential unalignments between LGMs, we have also integrated a deformable patch strategy, which involves both translation and rotation, as illustrated in **Figure 7b**. This approach enables LCM to capture a wider range of local geometry variations, allowing for robustness in rotation invariance. Specifically, the deformable patch strategy allows the LCM to flexibly adjust the spatial orientation and shape of the local surface patches to better match the geometry of the target object. For the translation patch strategy, we first sampled its local surface patches of the LGM with 1-ring neighborhood, where consists of six 1-hop neighbor vertices. Then, we slid the patches in six different directions to capture more information about the local geometry. Using the translation patch strategy, we obtained six LCMs for each pair of LGMs. As for the rotation patch strategy, we rotated each local surface patch with the sampled vertex at its center by ± 60⁰. We then obtained two LCMs for each pair of LGMs. We then maximized these LCMs and selected the final LCM for the identification procedure. This approach enabled us to efficiently capture and compare the local geometry of the infant cortical surface at each sampled vertex than vanilla LCM.

#### Vertex-wise LGM-based identification test

After computing all LCMs iteratively for all subjects in ***T***, we concatenated the ***n*** LCMs and formed a local correlation matrix, where each column represents an LCM of a pair of LGMs. Then, we can obtain the identity of the given subject by comparing the ***n*** LCMs. Then we leveraged a vertex-based identification solution, which predicts the identity for each sampled vertex by finding the individual with the highest similarity value at this vertex. Finally, through a majority voting across all sampled vertices, the identity presenting the highest frequency among all vertices was considered as the final predicted identity.

### Functional connectome-based identification test

The atlas-based individual infant identification test was performed across paired scans consisting of one target set, 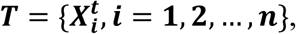 and one base set, 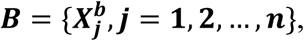 where 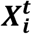 and 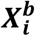 are all flattened vectors of the upper triangle of the whole-brain connectivity matrix and the subscripts ***i*** and ***j*** denote the subject index. With the requirement that 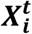 and 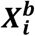 are acquired from the ***i***-th subject at different ages, each subject has two longitudinal fMRI scans from the two different sessions. In the process of identification, one target matrix, 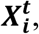 was selected from the target set ***T***, and the Pearson’s correlation coefficient was used to measure the similarity between 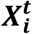 and all matrices in base set ***B***. The corresponding similarity matrix is defined as follows: 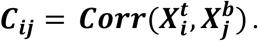 The identity in ***B*** having the largest similarity with 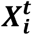 was regarded as the predicted identity of 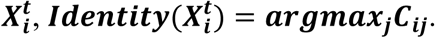 This procedure is shown in **Figure 7c**.

### Quantifying vertex-wise contributions to identification

Individual differences in functional connectivity were spatially inhomogeneous [1, 30]. To ascertain vertex-wise contribution to the individual identification process, we used two measures: uniqueness [30, 31] and differential power (DP) [1, 57]. LCMs showing high uniqueness values are shown consistent within same individual but varied widely among individuals, while those with high DP values are though to make a large contribution to individual identification.

#### Uniqueness

To identify the specific brain loci that manifest individual uniqueness and to assess the performance of FC fingerprint, we adopted the measure of uniqueness [30, 31], which qualifies the spatial distribution of inter-subject variability and provides new insights of individual differences. The uniqueness is calculated with averaged intra-individual local correlation maps ***C****_intra_* and averaged inter-individual local correlation maps ***C****_inter_*. For each dataset, the averaged intra-individual correlation map was derived from the same individual from different MRI sessions, 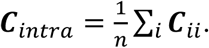 The average inter-individual correlation map was derived from different pairs of individuals from different MRI session, 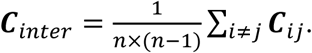 To estimate the uniqueness map, the intra-individual correlation map ***C****_intra_* and inter-individual correlation map ***C****_inter_* were subtracted from a vector of all ones, 𝕝, namely intra-individual variance and inter-individual variance. Then the intra-individual variance was regressed out from inter-individual variance using ordinary least squares regression. The residual map between regressed intra-individual variance and inter-individual variance was considered as the uniqueness map (𝕌). This process was as followed: 𝕌 = (𝕝 – ***C****_inter_*) − 𝛼(𝕝 – ***C****_intra_*) − *β*, where *α* and *β* are parameters determined via ordinary least-squares. Hence, the uniqueness map ***U*** would exhibit the highest values for measures that demonstrated substantial variability among individuals and could be consistent across multiple sessions within the same individual. The uniqueness maps derived from each dataset were then averaged for analysis.

#### Differential power

Given vertex *v* in LCM, if the vertex positively contributes to the identification of the individual, it should satisfy ***C***_***ii***_(*v*) > ***C***_***ij***_(*v*) for the identification process from session base to session target and ***C***_***ii***_(*v*) > ***C***_***ji***_(*v*) from session target to session base. The DP for vertex at the individual level is calculated as: 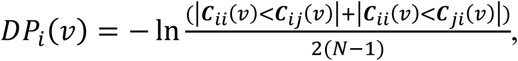 where |***C***_***ii***_(*v*) < ***C***_***ij***_(*v*)| and |***C***_***ii***_(*v*) < ***C***_***ji***_(*v*)| denote the total number of times that ***C***_***ii***_(*v*) < ***C***_***ij***_(*v*) and ***C***_***ii***_(*v*) < ***C***_***ji***_(*v*) across all other individual *j*, respectively. The vertex-wise DP for the infant population is calculated as: 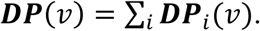 The larger the DP value is, the greater the contribution of this vertex to individual identification.

### Cognitive score prediction

To investigate the association between individual variability in local gradient-based functional connectivity fingerprints and infant cognitive development, we evaluated the predictive capacity of local gradient-based functional connectivity patterns for cognitive performance. Cognitive ability was assessed using the Early Learning Composite (ELC) score derived from the Mullen Early Learning Scale Assessment [53, 54], which aggregates scores from four domains: Fine Motor (FM), Visual Reception (VR), Receptive Language (RL), and Expressive Language (EL).

We employed a widely adopted machine learning model, i.e., the random forest (RF) to predict the ELC scores. The performance of our LGM with RF was evaluated through 10 iterations of 10-fold cross-validation, maintaining strict separation between training and testing sets to ensure data independence. In each iteration, during the training phase, we first conduct feature selection to select the most discriminative vertices and remove redundancy. Given that the local gradient features are sensitive to detecting localized abrupt transitions in resting-state functional connectivity patterns across the cortical surface, we extracted the local gradients of vertices located on boundary between brain regions as the representative features. A local gradient features-derived infant functional parcellation was employed at this step. Moreover, considering the potential unalignment between the parcellation and individual local gradient features, we further expand the boundary on the parcellation by dilating it three times to increase robustness. Then, the local gradient features of vertices within the same boundary between two brain regions were averaged and assigned as the feature of the specific boundary. Next, by correlating the features of each boundary with cognitive performance, the boundaries with significant correlation (p < 0.01) in the training set were identified and further categorized into two distinct feature sets based on their correlation directionality (positive or negative correlation with ELC scores). Later, we used the sum of these two distinct feature sets as a two-dimensional feature to predict the ELC scores with the random forest model. In the RF-based analysis, the number of ensemble trees equals 20, and the minimum number of observations per tree leaf equals five. During the testing phase, the fitted random forest model was used to predict the ELC scores directly.

To compute the differential contributions of functional networks to cognitive prediction. We computed the number of vertices in the selected boundaries, normalized by the total vertex count of all boundaries within each network to account for the heterogeneity of network size and boundary number. Specifically, for the boundaries located between the networks, they were counted for both networks when computing contributions.

Model performance was evaluated by averaging predictions across 10 iterations of 10-fold cross-validation. Predictive accuracy was assessed through Pearson’ correlation coefficient between predicted and observed scores.

## Acknowledgements

This work was supported in part by NIH grants (MH123202, ES033518, AG075582, NS128534, and NS135574). This work also utilizes approaches developed by an NIH grant (1U01MH110274) and the efforts of the UNC/UMN Baby Connectome Project Consortium.

## Author Contributions

X.Y. and G.L. designed research; X.Y., J.C., D.H., Z.W., W.L., L.W., and G.L. performed research and analyzed data; X.Y., J.C., D.H., Z.W., W.L., L.W., and G.L. wrote the paper.

## Ethic Declaration

The authors declare that they have no conflict of interest.

## Data Availability

All MRI data analyzed during this study are from the UNC/UMN Baby Connectome Project (https://nda.nih.gov/edit_collection.html?id=2848), which is publicly available in NIMH Data Archive.

## Code Availability

This study used openly available software and codes. The computation and analysis codes are available at https://github.com/BRAIN-Lab-UNC/LocalGradFingerprint.

## Notes

### Competing Interest Statement

The authors have declared no competing interest.

## Reference

1. Emily S Finn, Xilin Shen, Dustin Scheinost, Monica D Rosenberg, Jessica Huang, Marvin M Chun, Xenophon Papademetris, and R Todd Constable. Functional connectome fingerprinting: identifying individuals using patterns of brain connectivity. Nature neuroscience, 18(11):1664–1671, 2015.

2. Corey Horien, Xilin Shen, Dustin Scheinost, and R Todd Constable. The individual functional connectome is unique and stable over months to years. Neuroimage, 189:676–687, 2019.

3. Oscar Miranda-Dominguez, Eric Feczko, David S Grayson, Hasse Walum, Joel T Nigg, and Damien A Fair. Heritability of the human connectome: A connectotyping study. Network Neuroscience, 2(02):175–199, 2018.

4. Damion V Demeter, Laura E Engelhardt, Remington Mallett, Evan M Gordon, Tehila Nugiel, K Paige Harden, Elliot M Tucker-Drob, Jarrod A Lewis-Peacock, and Jessica A Church. Functional connectivity fingerprints at rest are similar across youths and adults and vary with genetic similarity. Iscience, 23(1), 2020.

5. Tobias Kaufmann, Dag Alnæs, Nhat Trung Doan, Christine Lycke Brandt, Ole A Andreassen, and Lars T Westlye. Delayed stabilization and individualization in connectome development are related to psychiatric disorders. Nature neuroscience, 20(4):513–515, 2017.

6. Maria Jalbrzikowski, Fuchen Liu, William Foran, Lambertus Klei, Finnegan J Calabro, Kathryn Roeder, Bernie Devlin, and Beatriz Luna. Functional connectome fingerprinting accuracy in youths and adults is similar when examined on the same day and 1.5-years apart. Human brain mapping, 41(15):4187–4199, 2020.

7. John H Gilmore, Rebecca C Knickmeyer, and Wei Gao. Imaging structural and functional brain development in early childhood. Nature Reviews Neuroscience, 19(3):123–137, 2018.

8. Gang Li, Jingxin Nie, Li Wang, Feng Shi, Amanda E Lyall, Weili Lin, John H Gilmore, and Dinggang Shen. Mapping longitudinal hemispheric structural asymmetries of the human cerebral cortex from birth to 2 years of age. Cerebral cortex, 24(5):1289–1300, 2014.

9. Sarah J Paterson, Sabine Heim, Jennifer Thomas Friedman, Naseem Choudhury, and April A Benasich. Development of structure and function in the infant brain: Implications for cognition, language and social behaviour. Neuroscience & Biobehavioral Reviews, 30(8):1087–1105, 2006.

10. Alice M Graham, Jennifer H Pfeifer, Philip A Fisher, Weili Lin, Wei Gao, and Damien A Fair. The potential of infant fmri research and the study of early life stress as a promising exemplar. Developmental cognitive neuroscience, 12:12–39, 2015.

11. Han Zhang, Dinggang Shen, and Weili Lin. Resting-state functional mri studies on infant brains: a decade of gap-filling efforts. Neuroimage, 185:664–684, 2019.

12. Wei Gao, Sarael Alcauter, J Keith Smith, John H Gilmore, and Weili Lin. Development of human brain cortical network architecture during infancy. Brain Structure and Function, 220:1173–1186, 2015.

13. Evan M Gordon, Timothy O Laumann, Babatunde Adeyemo, Jeremy F Huckins, William M Kelley, and Steven E Petersen. Generation and evaluation of a cortical area parcellation from resting-state correlations. Cerebral cortex, 26(1):288–303, 2016.

14. Dan Hu, Fan Wang, Han Zhang, Zhengwang Wu, Zhen Zhou, Guoshi Li, Li Wang, Weili Lin, Gang Li, UNC/UMN Baby Connectome Project Consortium, et al. Existence of functional connectome fingerprint during infancy and its stability over months. Journal of Neuroscience, 42(3):377–389, 2022.

15. Fan Wang, Han Zhang, Zhengwang Wu, Dan Hu, Zhen Zhou, Jessica B Girault, Li Wang, Weili Lin, and Gang Li. Fine-grained functional parcellation maps of the infant cerebral cortex. elife, 12:e75401, 2023.

16. Ru Kong, Qing Yang, Evan Gordon, Aihuiping Xue, Xiaoxuan Yan, Csaba Orban, Xi-Nian Zuo, Nathan Spreng, Tian Ge, Avram Holmes, et al. Individual-specific areal-level parcellations improve functional connectivity prediction of behavior. Cerebral cortex, 31(10):4477–4500, 2021.

17. Boris C Bernhardt, Jonathan Smallwood, Shella Keilholz, and Daniel S Margulies. Gradients in brain organization, 2022.

18. Alexander Schaefer, Ru Kong, Evan M Gordon, Timothy O Laumann, Xi-Nian Zuo, Avram J Holmes, Simon B Eickhoff, and BT Thomas Yeo. Local-global parcellation of the human cerebral cortex from intrinsic functional connectivity mri. Cerebral cortex, 28(9):3095–3114, 2018.

19. Gagan S Wig, Timothy O Laumann, and Steven E Petersen. An approach for parcellating human cortical areas using resting-state correlations. Neuroimage, 93:276–291, 2014.

20. Daniel S Margulies, Satrajit S Ghosh, Alexandros Goulas, Marcel Falkiewicz, Julia M Huntenburg, Georg Langs, Gleb Bezgin, Simon B Eickhoff, F Xavier Castellanos, Michael Petrides, et al. Situating the default-mode network along a principal gradient of macroscale cortical organization. Proceedings of the National Academy of Sciences, 113(44):12574– 12579, 2016.

21. Ye Tian, Daniel S Margulies, Michael Breakspear, and Andrew Zalesky. Topographic organization of the human subcortex unveiled with functional connectivity gradients. Nature neuroscience, 23(11):1421–1432, 2020.

22. Annchen R Knodt, Maxwell L Elliott, Ethan T Whitman, Alex Winn, Angela Addae, David Ireland, Richie Poulton, Sandhya Ramrakha, Avshalom Caspi, Terrie E Moffitt, et al. Test– retest reliability and predictive utility of a macroscale principal functional connectivity gradient. Human Brain Mapping, 44(18):6399–6417, 2023.

23. Seok-Jun Hong, Ting Xu, Aki Nikolaidis, Jonathan Smallwood, Daniel S Margulies, Boris Bernhardt, Joshua Vogelstein, and Michael P Milham. Toward a connectivity gradient-based framework for reproducible biomarker discovery. NeuroImage, 223:117322, 2020.

24. Zhiyao Gao, Li Zheng, Katya Krieger-Redwood, Ajay Halai, Daniel S Margulies, Jonathan Smallwood, and Elizabeth Jefferies. Flexing the principal gradient of the cerebral cortex to suit changing semantic task demands. Elife, 11:e80368, 2022.

25. Xuyun Wen, Han Zhang, Gang Li, Mingxia Liu, Weiyan Yin, Weili Lin, Jun Zhang, and Dinggang Shen. First-year development of modules and hubs in infant brain functional networks. Neuroimage, 185:222–235, 2019.

26. Wei Gao, Sarael Alcauter, Amanda Elton, Carlos R Hernandez-Castillo, J Keith Smith, Juanita Ramirez, and Weili Lin. Functional network development during the first year: relative sequence and socioeconomic correlations. Cerebral cortex, 25(9):2919–2928, 2015.

27. Arnaud Messé. Parcellation influence on the connectivity-based structure–function relationship in the human brain. Human Brain Mapping, 41(5):1167–1180, 2020.

28. Brittany R Howell, Martin A Styner, Wei Gao, Pew-Thian Yap, Li Wang, Kristine Baluyot, Essa Yacoub, Geng Chen, Taylor Potts, Andrew Salzwedel, et al. The unc/umn baby connectome project (bcp): An overview of the study design and protocol development. NeuroImage, 185:891–905, 2019.

29. Matthew F Glasser, Timothy S Coalson, Emma C Robinson, Carl D Hacker, John Harwell, Essa Yacoub, Kamil Ugurbil, Jesper Andersson, Christian F Beckmann, Mark Jenkinson, et al. A multi-modal parcellation of human cerebral cortex. Nature, 536(7615):171–178, 2016.

30. Ye Tian, BT Thomas Yeo, Vanessa Cropley, Andrew Zalesky, et al. High-resolution connectomic fingerprints: Mapping neural identity and behavior. NeuroImage, 229:117695, 2021.

31. Sophia Mueller, Danhong Wang, Michael D Fox, BT Thomas Yeo, Jorge Sepulcre, Mert R Sabuncu, Rebecca Shafee, Jie Lu, and Hesheng Liu. Individual variability in functional connectivity architecture of the human brain. Neuron, 77(3):586–595, 2013.

32. Jin Liu, Xuhong Liao, Mingrui Xia, and Yong He. Chronnectome fingerprinting: Identifying individuals and predicting higher cognitive functions using dynamic brain connectivity patterns. Human brain mapping, 39(2):902–915, 2018.

33. Judit Ciarrusta, Daan Christiaens, Sean P Fitzgibbon, Ralica Dimitrova, Jana Hutter, Emer Hughes, Eugene Duff, Anthony N Price, Lucilio Cordero-Grande, J-Donald Tournier, et al. The developing brain structural and functional connectome fingerprint. Developmental Cognitive Neuroscience, 55:101117, 2022.

34. Tamara Vanderwal, Jeffrey Eilbott, Clare Kelly, Simon R Frew, Todd S Woodward, Michael P Milham, and F Xavier Castellanos. Stability and similarity of the pediatric connectome as developmental measures. NeuroImage, 226:117537, 2021.

35. Daouia I Larabi, Martin Gell, Enrico Amico, Simon B Eickhoff, and Kaustubh R Patil. Highly accurate local functional fingerprints and their stability. bioRxiv, 8(03):454862, 2021.

36. Yasuo Mori, Jun Miyata, Masanori Isobe, Shuraku Son, Yujiro Yoshihara, Toshihiko Aso, Takanori Kouchiyama, Toshiya Murai, and Hidehiko Takahashi. Effect of phase-encoding direction on group analysis of resting-state functional magnetic resonance imaging. Psychiatry and clinical neurosciences, 72(9):683–691, 2018.

37. Dafnis Batalle, A David Edwards, and Jonathan O’Muircheartaigh. Annual research review: not just a small adult brain: understanding later neurodevelopment through imaging the neonatal brain. Journal of Child Psychology and Psychiatry, 59(4):350–371, 2018.

38. Andreas Burkhalter. Development of forward and feedback connections between areas v1 and v2 of human visual cortex. Cerebral cortex, 3(5):476–487, 1993.

39. Wenjian Gao, Ziyi Huang, Wenfei Ou, Xiaoqian Tang, Wanying Lv, and Jingxin Nie. Functional individual variability development of the neonatal brain. Brain Structure and Function, 227(6):2181–2190, 2022.

40. Qiushi Wang, Yuehua Xu, Tengda Zhao, Zhilei Xu, Yong He, and Xuhong Liao. Individual uniqueness in the neonatal functional connectome. Cerebral Cortex, 31(8):3701–3712, 2021.

41. Peter R Huttenlocher and Arun S Dabholkar. Regional differences in synaptogenesis in human cerebral cortex. Journal of comparative Neurology, 387(2):167–178, 1997.

42. Miao Cao, Hao Huang, and Yong He. Developmental connectomics from infancy through early childhood. Trends in neurosciences, 40(8):494–506, 2017.

43. Sophia Stoecklein, Anne Hilgendorff, Meiling Li, Kai Förster, Andreas W Flemmer, Franziska Galiè, Stephan Wunderlich, Danhong Wang, Sophie Stein, Harald Ehrhardt, et al. Variable functional connectivity architecture of the preterm human brain: impact of developmental cortical expansion and maturation. Proceedings of the National Academy of Sciences, 117(2):1201–1206, 2020.

44. Yuehua Xu, Miao Cao, Xuhong Liao, Mingrui Xia, Xindi Wang, Tina Jeon, Minhui Ouyang, Lina Chalak, Nancy Rollins, Hao Huang, et al. Development and emergence of individual variability in the functional connectivity architecture of the preterm human brain. Cerebral Cortex, 29(10):4208–4222, 2019.

45. Michael W Cole, Tal Yarkoni, Grega Repovš, Alan Anticevic, and Todd S Braver. Global connectivity of prefrontal cortex predicts cognitive control and intelligence. Journal of Neuroscience, 32(26):8988–8999, 2012.

46. Valentina Doria, Christian F Beckmann, Tomoki Arichi, Nazakat Merchant, Michela Groppo, Federico E Turkheimer, Serena J Counsell, Maria Murgasova, Paul Aljabar, Rita G Nunes, et al. Emergence of resting state networks in the preterm human brain. Proceedings of the National Academy of Sciences, 107(46):20015–20020, 2010.

47. Christopher D Smyser and Jeffrey J Neil. Use of resting-state functional mri to study brain development and injury in neonates. In Seminars in perinatology, volume 39, pages 130–140. Elsevier, 2015.

48. Miao Cao, Yong He, Zhengjia Dai, Xuhong Liao, Tina Jeon, Minhui Ouyang, Lina Chalak, Yanchao Bi, Nancy Rollins, Qi Dong, et al. Early development of functional network segregation revealed by connectomic analysis of the preterm human brain. Cerebral cortex, 27(3):1949–1963, 2017.

49. Ru Kong, Yan Rui Tan, Naren Wulan, Leon Qi Rong Ooi, Seyedeh-Rezvan Farahibozorg, Samuel Harrison, Janine D Bijsterbosch, Boris C Bernhardt, Simon Eickhoff, and BT Thomas Yeo. Comparison between gradients and parcellations for functional connectivity prediction of behavior. NeuroImage, 273:120044, 2023.

50. Timothy O Laumann, Evan M Gordon, Babatunde Adeyemo, Abraham Z Snyder, Sung Jun Joo, Mei-Yen Chen, Adrian W Gilmore, Kathleen B McDermott, Steven M Nelson, Nico UF Dosenbach, et al. Functional system and areal organization of a highly sampled individual human brain. Neuron, 87(3):657–670, 2015.

51. Jesse A Brown, Alex J Lee, Lorenzo Pasquini, and William W Seeley. A dynamic gradient architecture generates brain activity states. NeuroImage, 261:119526, 2022.

52. Richard AI Bethlehem, Casey Paquola, Jakob Seidlitz, Lisa Ronan, Boris Bernhardt, Kamen A Tsvetanov, Cam-CAN Consortium, et al. Dispersion of functional gradients across the adult lifespan. Neuroimage, 222:117299, 2020.

53. Eileen M Mullen et al. Mullen scales of early learning. AGS Circle Pines, MN, 1995.

54. Neta Yitzhak, Ayelet Harel, Maya Yaari, Edwa Friedlander, and Nurit Yirmiya. The mullen scales of early learning: ceiling effects among preschool children. European Journal of Developmental Psychology, 13:1–14, 11 2015.

55. Gang Li, Li Wang, Pew-Thian Yap, Fan Wang, Zhengwang Wu, Yu Meng, Pei Dong, Jaeil Kim, Feng Shi, Islem Rekik, et al. Computational neuroanatomy of baby brains: A review. NeuroImage, 185:906–925, 2019.

56. Zhengwang Wu, Li Wang, Weili Lin, John H Gilmore, Gang Li, and Dinggang Shen. Construction of 4d infant cortical surface atlases with sharp folding patterns via spherical patch-based group-wise sparse representation. Human brain mapping, 40(13):3860–3880, 2019.

57. Gang Li, Li Wang, Feng Shi, John H Gilmore, Weili Lin, and Dinggang Shen. Construction of 4d high-definition cortical surface atlases of infants: Methods and applications. Medical image analysis, 25(1):22–36, 2015.

58. Li Wang, Zhengwang Wu, Liangjun Chen, Yue Sun, Weili Lin, and Gang Li. Ibeat v2. 0: a multisite-applicable, deep learning-based pipeline for infant cerebral cortical surface reconstruction. Nature protocols, 18(5):1488–1509, 2023.

59. Stephen M Smith, Mark Jenkinson, Mark W Woolrich, Christian F Beckmann, Timothy EJ Behrens, Heidi Johansen-Berg, Peter R Bannister, Marilena De Luca, Ivana Drobnjak, David E Flitney, et al. Advances in functional and structural mr image analysis and implementation as fsl. Neuroimage, 23:S208–S219, 2004.

60. Mark Jenkinson, Peter Bannister, Michael Brady, and Stephen Smith. Improved optimization for the robust and accurate linear registration and motion correction of brain images. Neuroimage, 17(2):825–841, 2002.

61. Mark W Woolrich, Saad Jbabdi, Brian Patenaude, Michael Chappell, Salima Makni, Timothy Behrens, Christian Beckmann, Mark Jenkinson, and Stephen M Smith. Bayesian analysis of neuroimaging data in fsl. Neuroimage, 45(1):S173–S186, 2009.

62. Qian Zhang, Li Wang, Xiaopeng Zong, Weili Lin, Gang Li, and Dinggang Shen. Frnet: Flattened residual network for infant mri skull stripping. In 2019 IEEE 16^th^ International Symposium on Biomedical Imaging (ISBI 2019), pages 999–1002. IEEE, 2019.

63. John G Sled, Alex P Zijdenbos, and Alan C Evans. A nonparametric method for automatic correction of intensity nonuniformity in mri data. IEEE transactions on medical imaging, 17(1):87–97, 1998.

64. Li Wang, Gang Li, Feng Shi, Xiaohuan Cao, Chunfeng Lian, Dong Nie, Mingxia Liu, Han Zhang, Guannan Li, Zhengwang Wu, et al. Volume-based analysis of 6-month-old infant brain mri for autism biomarker identification and early diagnosis. In Medical Image Computing and Computer Assisted Intervention–MICCAI 2018: 21st International Conference, Granada, Spain, September 16-20, 2018, Proceedings, Part III 11, pages 411–419. Springer, 2018.

65. Zhen Zhou, Han Zhang, Li-Ming Hsu, Weili Lin, Gang Pan, Dinggang Shen, and UNC/UMN Baby Connectome Project Consortium. Multi-layer temporal network analysis reveals increasing temporal reachability and spreadability in the first two years of life. In Medical Image Computing and Computer Assisted Intervention–MICCAI 2019: 22nd International Conference, Shenzhen, China, October 13–17, 2019, Proceedings, Part III 22, pages 665–672. Springer, 2019.

66. Douglas N Greve and Bruce Fischl. Accurate and robust brain image alignment using boundary-based registration. Neuroimage, 48(1):63–72, 2009.

67. Ludovica Griffanti, Gholamreza Salimi-Khorshidi, Christian F Beckmann, Edward J Auerbach, Gwenaëlle Douaud, Claire E Sexton, Enikő Zsoldos, Klaus P Ebmeier, Nicola Filippini, Clare E Mackay, et al. Ica-based artefact removal and accelerated fmri acquisition for improved resting state network imaging. Neuroimage, 95:232–247, 2014.

68. Christian F Beckmann and Stephen M Smith. Probabilistic independent component analysis for functional magnetic resonance imaging. IEEE transactions on medical imaging, 23(2):137– 152, 2004.

69. Tae-Eui Kam, Xuyun Wen, Bing Jin, Zhicheng Jiao, Li-Ming Hsu, Zhen Zhou, Yujie Liu, Koji Yamashita, Sheng-Che Hung, Weili Lin, et al. A deep learning framework for noise component detection from resting-state functional mri. In Medical Image Computing and Computer Assisted Intervention–MICCAI 2019: 22nd International Conference, Shenzhen, China, October 13–17, 2019, Proceedings, Part III 22, pages 754–762. Springer, 2019.

70. Gang Li, Jingxin Nie, Guorong Wu, Yaping Wang, Dinggang Shen, Alzheimer’s Disease Neuroimaging Initiative, et al. Consistent reconstruction of cortical surfaces from longitudinal brain mr images. Neuroimage, 59(4):3805–3820, 2012.

71. Gang Li, Jingxin Nie, Li Wang, Feng Shi, John H Gilmore, Weili Lin, and Dinggang Shen. Measuring the dynamic longitudinal cortex development in infants by reconstruction of temporally consistent cortical surfaces. Neuroimage, 90:266–279, 2014.

72. Bruce Fischl, Martin I Sereno, and Anders M Dale. Cortical surface-based analysis: Ii: inflation, flattening, and a surface-based coordinate system. Neuroimage, 9(2):195–207, 1999.

73. BT Thomas Yeo, Mert R Sabuncu, Tom Vercauteren, Nicholas Ayache, Bruce Fischl, and Polina Golland. Spherical demons: fast diffeomorphic landmark-free surface registration. IEEE transactions on medical imaging, 29(3):650–668, 2009.

74. Matthew F Glasser, Stamatios N Sotiropoulos, J Anthony Wilson, Timothy S Coalson, Bruce Fischl, Jesper L Andersson, Junqian Xu, Saad Jbabdi, Matthew Webster, Jonathan R Polimeni, et al. The minimal preprocessing pipelines for the human connectome project. Neuroimage, 80:105–124, 2013.

